# Key Factors Governing Initial Stages of Lipid Droplet Formation

**DOI:** 10.1101/2021.11.12.468423

**Authors:** Siyoung Kim, Chenghan Li, Robert V. Farese, Tobias C. Walther, Gregory A. Voth

## Abstract

Lipid droplets (LDs) are neutral lipid storage organelles surrounded by a phospholipid (PL) monolayer. LD biogenesis from the endoplasmic reticulum (ER) is driven by phase separation of neutral lipids, overcoming surface tension and membrane deformation. However, the core biophysics of the initial steps of LD formation remain relatively poorly understood. Here, we use a tunable, phenomenological coarse-grained (CG) model to study triacylglycerol (TG) nucleation in a bilayer membrane. We show that PL rigidity has a strong influence on TG lensing and membrane remodeling: When membrane rigidity increases, TG clusters remain more planar with high anisotropy but a minor degree of phase nucleation. This finding is confirmed by free energy sampling simulations that calculate the potential of mean force (PMF) as a function of the degree of nucleation and anisotropy. We also show that asymmetric tension, controlled by the number of PLs on each membrane leaflet, determines the budding direction. A TG lens buds in the direction of the monolayer containing excess PLs to allow for better PL coverage of TG, consistent with reported experiments. Finally, two governing mechanisms of the LD growth, Ostwald ripening and merging, are observed. Taken together, this study characterizes the interplay between two thermodynamic quantities during the initial LD phases, the TG bulk free energy and membrane remodeling energy.

## INTRODUCTION

Lipid droplets (LDs) are ubiquitous organelles that store lipids. LDs are considered an oil-in-water emulsion in a cell with their core consisting of neutral lipids such as triacylglycerol (TG) or sterol esters, surrounded by a phospholipid (PL) monolayer.^1–4^ During LD formation from the endoplasmic reticulum (ER), cells package neutral lipids with a PL monolayer. Lipid droplet assembly complexes (LDACs), and in particular the ER protein seipin, determine where LDs form and facilitate the process.^5^

Biophysically, LD emergence can be considered in the context of classical nucleation theory.^6^ The driving force of TG nucleation is the bulk energy of TG, stemming from the hydrophobic interactions of TG’s three acyl chains and polar interactions between TG’s glycerol moieties. As LDs grow they inflict a raising energy penalty due to surface tension (~1 mN/m), proportional to the LD surface area.^7^ In addition, TG lensing in the ER membrane leads to deformation of the membrane. The energy penalty due to membrane deformation is more dominant than the surface tension term during the initial phases of LD formation when the phase boundary between the forming LD and cytoplasm is small. However, as the LD surface expands with LD growth, the surface tension term becomes dominant.^8^ Therefore, one may expect that an initial TG lens is flat to reduce membrane deformation, and it becomes more and more spherical to reduce the surface tension penalty as the LD grows. Such a process has been predicted in theory^9–10^ and shown in molecular dynamics (MD) simulations.^11^

Due to their limited time and length scales, studying LD biogenesis with all-atom (AA) MD simulations is not viable. For instance, TG and diacylglycerol (DAG) molecules do not nucleate in bilayers during 1 μs at the higher concentrations than the critical.^12^ Therefore, in this work, we study initial LD biogenesis with a tunable, coarse-grained (CG) PL model^13^ and a TG model derived from it. Using the CG model, we aim to understand the initial LD formation in the regime where the membrane deformation penalty is more significant than the surface tension penalty. We report the interplay between the shape of a TG blister and PL rigidity, Ostwald ripening, the tension-dependent budding behavior, and the calculation of TG nucleation potential of mean force (PMF).

## METHODS

### CG lipid model

An implicit solvent (solvent-free), phenomenological CG model for PL and TG was used (Fig. 1). Each PL and TG molecule consists of four CG beads. Despite being a linear model, it correctly represents the number of acyl chains that each molecule has and the relative effect of hydrophobic interactions. A PL molecule has two tail atoms while a TG molecule has three tail atoms in the CG model, consistent with the number of acyl chains in their chemical structures. No CG bead carried any charge. All CG pairs except bonded ones interact with each other with the following pair potential,

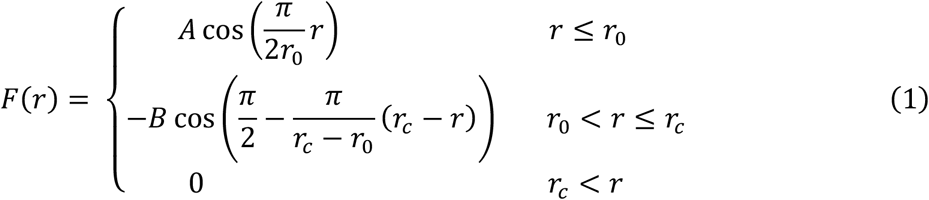

where *r*_*c*_ is 2*r*_0_.^13^ The repulsion is a sine-based soft-core repulsion and much softer than a hard-core repulsion such as Lennard-Jones, which allows a larger integration timestep. In addition, the absence of electrostatic interactions, the relatively short cutoff distance (1.5 nm) of nonbonded interactions (considering the removal of charges), and the low CG resolution enable fast and efficient calculations to access large length and time scales. The repulsion parameter, *A*, was chosen as 25 *k*_B_*T*, consistent with the original PL model.^13^ The attraction occurs between the following pairs of atom types: PGL-PGL, T-T, TGL-TGL, and TGL-T. The attraction parameter, *B*, was set to 1 *k*_B_*T* except for the TGL-TGL pair, which was chosen as 1.1 k_B_T. The higher attraction in the TGL-TGL pair was motivated by the previous paper that shows a sharper radial distribution function between the TG glycerol moieties than any other PL or TG pairs from the mapped atomistic trajectories.^12^ The other pair is purely repulsive by setting the attraction parameter to zero. The parameter *r*_0_ was set to 7.5 Å except for the interaction between the head and head groups, which was set to 4.5 Å. Both potential and force become zero at the cutoff distance (*r*_*c*_), therefore there is no need for a switching or shifting function. The bonded interaction is harmonic with an equilibrium distance of 7.5 Å and a spring constant of 25 *k*_B_*T*/Å^2^. Each PL molecule has two harmonic angles with an equilibrium angle of 180° and a spring constant of 0.5 *k*_B_*T* (soft PL) or 2 *k*_B_*T* (stiff PL). The TG molecules do not have angle potentials. The mass of each CG bead was chosen as 200 g/mol because a molecular weight of PL or TG is in a range of 600 g/mol – 900 g/mol. We note that this is a purely phenomenological CG model, and therefore the quantities calculated here are not directly related to the underlying all-atom system.

**FIGURE 1.**
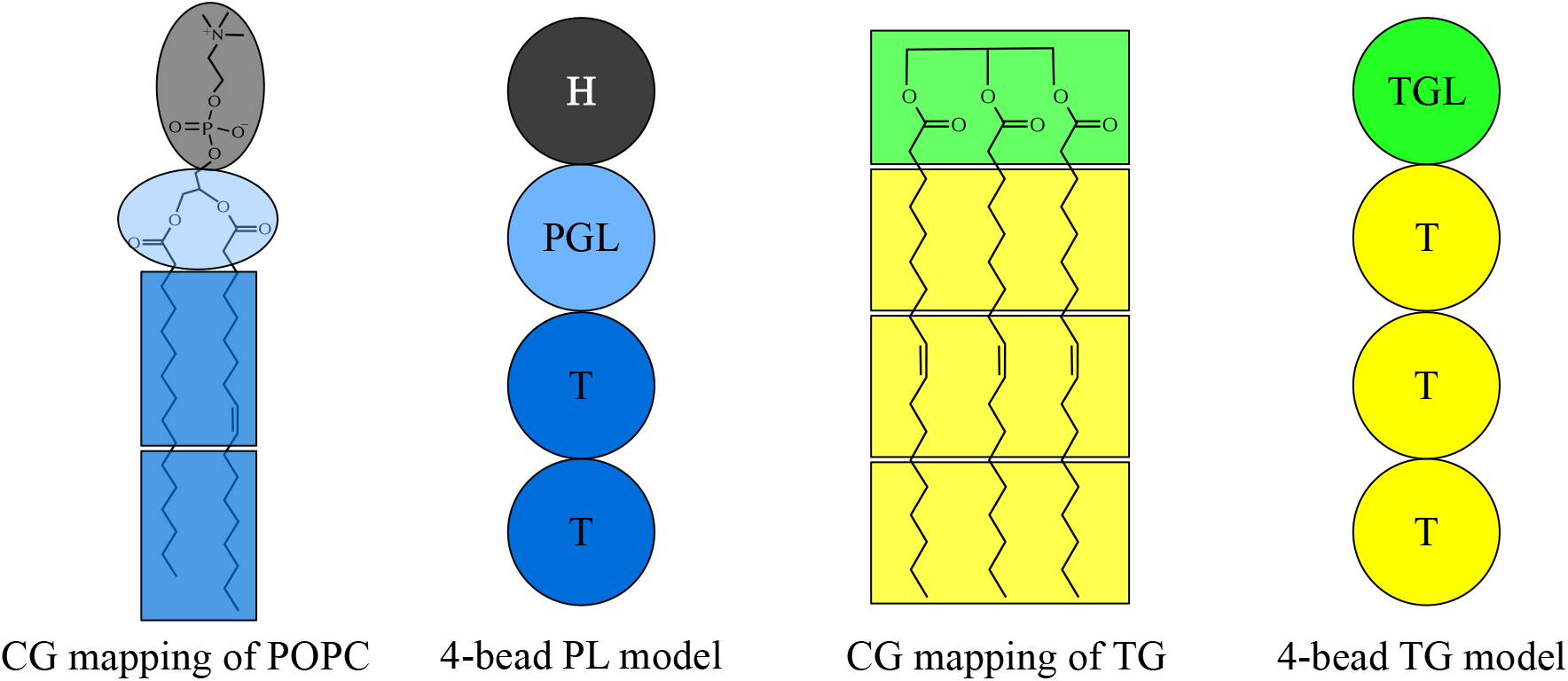
Schematic representation of the mapping of PL and TG.

### All-Atom simulations

A 1-palmitoyl-2-oleoyl-sn-glycero-3-phosphocholine (POPC) bilayer membrane, consisting of 64 molecules in each leaflet, was constructed using the CHARMM-GUI membrane builder.^14–15^ The production run was conducted for 200 ns by GROMACS 2018^16^ with the CHARMM36 force field.^17^ The Lennard-Jones interaction was force-switched between 1.0 nm to 1.2 nm. Simulations were evolved with a 2-fs timestep. The particle mesh Ewald algorithm^18^ was used to evaluate the long-range electrostatic interactions with a real distance cutoff of 1.2 nm. Any bond involving a hydrogen atom was constrained using the LINCS algorithm.^19^ The Nose-Hoover thermostat was used with a target temperature of 310 K and with a coupling time constant of 1 ps.^20–21^ Semi-isotropically coupled pressure was controlled with the Parrinello-Rahman barostat with a target pressure of 1 bar and with a compressibility of 4.5 × 10^−5^ bar^−1^ and a coupling time constant of 5 ps.^22^

### Coarse-grained simulations

The CG MD simulations were carried out using LAMMPS with tabulated CG potentials.^23^ Simulations were evolved with a 50-fs timestep. The Langevin thermostat was used with a target temperature of 310 K and with a coupling constant of 100 ps.^24^ In a flat bilayer simulation, the Nose-Hoover barostat with the Martyna-Tobias-Klein correction was used with a target pressure of 0 atm in the *XY* dimension and with a coupling constant of 250 ps.^22, 25–26^ The pressure in the *X* and *Y* dimensions was coupled. The cutoff distance of nonbonded interactions was set to 1.5 nm, where both the force and potential become zero. Biased simulations (described below) were performed with the external plugin PLUMED2.6.^27^ The initial structures of the CG simulations were prepared with the software MDAnalysis.^28^ Simulation details are provided in Table 1.

**Table 1.**
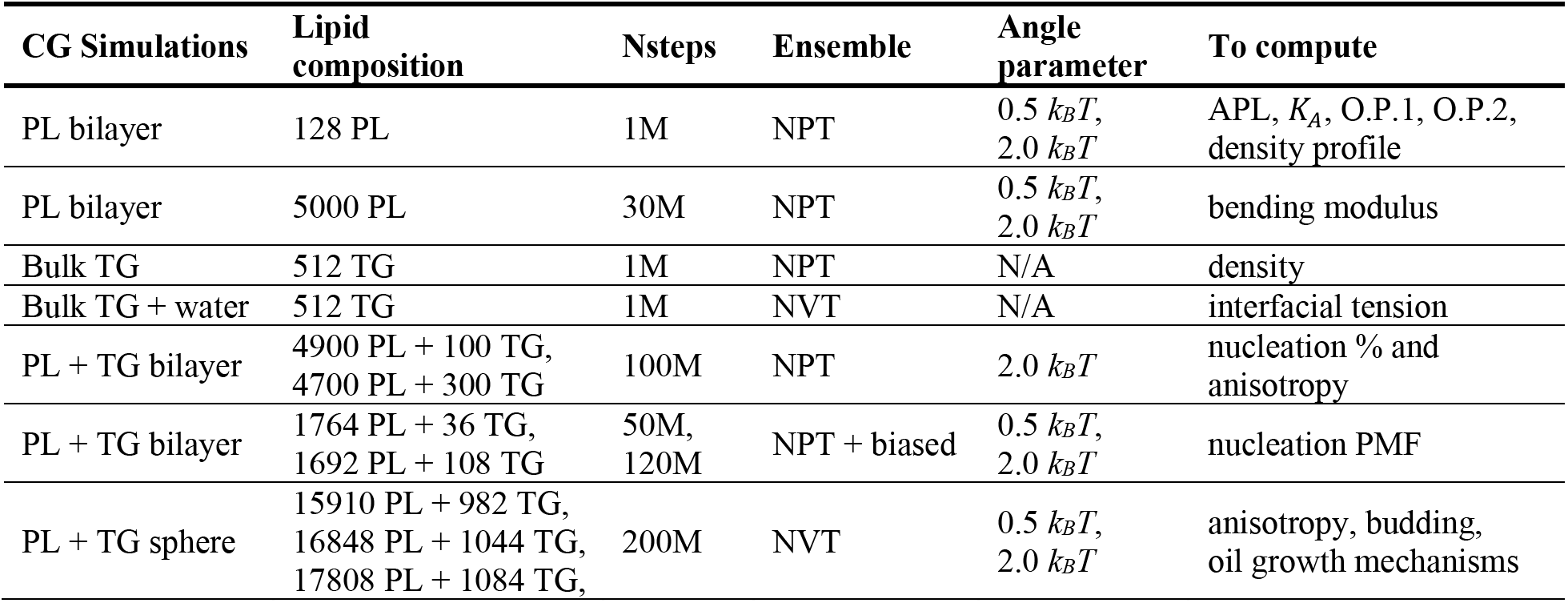
Description of CG simulations.

### Nucleation percentage and anisotropy

We mainly calculated two quantities in this study: the nucleation percentage and anisotropy. The former was defined as the number of TG molecules in the largest cluster divided by the total number of TG molecules in the system. If two TGL atoms are within 2 nm (chosen by inspection of CG trajectories), those two TG molecules are considered to be in one cluster.

Anisotropy (*k*) describes the shape of a TG lens and is computed with the following procedure: First, we identify the largest TG cluster. Second, we calculate the moment of inertia tensor of those TG molecules from the center of mass of the TG lens. Third, the moment of inertia tensor is diagonalized. Finally, we calculate the anisotropy as:

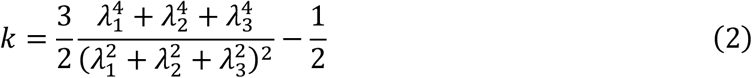

where *λ*_1_, *λ*_2_, and *λ*_3_ are the eigenvalues of the inertia tensor. Anisotropy ranges from 0 to 0.25, where 0 represents a sphere, and 0.25 does a plane.

### Area compressibility

Bilayer area compressibility was calculated using the following equation

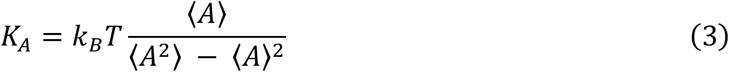

where *A* is the area of a membrane.

### Order parameters

The PL order parameters were calculated with the following equation, *S*_*CD*_ = 0.5 × |〈3 cos^2^ *θ* − 1〉|, where the angle (*θ*) is between the *Z* axis and the position vector of a tail atom to a glycerol group (PGL). If the positional vector is from the first tail atom, which is closer to the PGL atom, the calculated quantity is referred to as O.P.1, and if the vector is from the second tail atom, the quantity is referred to as O.P.2. The AA MD trajectory was first mapped with the mapping scheme illustrated in Fig. 1, followed by the calculation of the order parameters.

### TG nucleation PMF

Well-tempered metadynamics simulations^29^ were carried out to compute the TG nucleation PMF using the same procedure described in Ref. 30. The Gaussian hills were deposited every 500 steps at the height of 0.48 kcal/mol and a width of 10. A biasfactor of 50 was used. The biased collective variable was the sum of the coordination number of the TG glycerol atoms that were in the largest TG cluster. A switching function was defined to compute the coordination number,

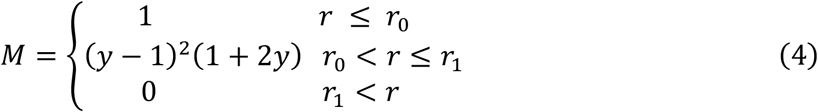

where 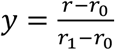, *r*_0_ = 1.95 *nm*, and *r*_1_ = 2.00 *nm*. The above switching function is plotted in Fig. 2. By defining the function that switches to 0 at 2.0 nm, this collective variable has a consistent cutoff distance with the nucleation percentage. The biased trajectories were reweighted with the following equation,

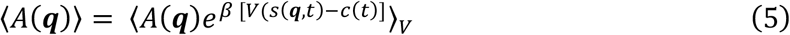

where *A* and ***q*** represent the property of the interest and the coordinates of atoms, respectively.^31–32^ The terms *V* and *s* represent the biasing potential and collective variable, respectively. The time-dependent constant was calculated as

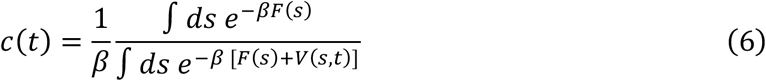

The final potential of mean force (PMF) was represented with the nucleation percentage and anisotropy. The PLUMED script that biased the simulations is included in the Supporting Information.

**FIGURE 2.**
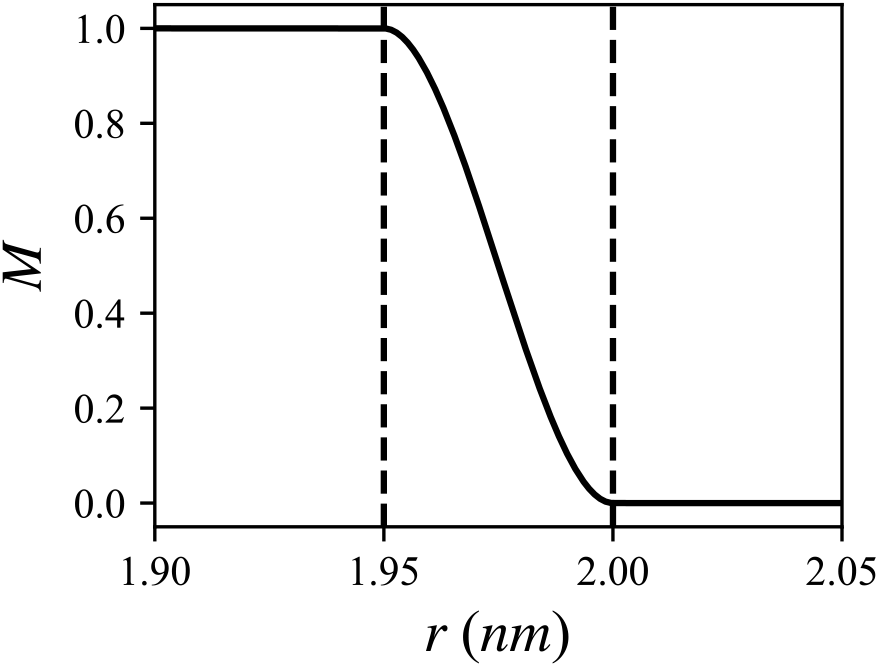
Switching function used in the calculation of the coordination number.

## RESULTS

### Physical properties of PL and TG

First, we characterized the physical properties of PL bilayers and bulk TG. We prepared a PL bilayer, consisting of 64 PL molecules in each leaflet, to compute the area per lipid (APL), area compressibility, order parameter (Table 2), and number density profile (Fig. 3). Two angle potential parameters, 0.5 *k*_B_*T* and 2.0 *k*_B_*T*, were tested. Both soft and stiff PLs had a liquid-disordered phase at 310 K. We also set up a larger PL bilayer, consisting of 2500 PL molecules in each leaflet, to calculate bending modulus. We first note how angle parameters modulate the physical properties of bilayers. As PL stiffness increased, PLs became more rigid, therefore reducing the APL and increasing the bending modulus, area compressibility, and order parameters (Table 2). This implies that the initial LD formation would be harder, and a TG blister would be more planar with a higher angle potential parameter, which will be discussed later. The same relation between PL rigidity and an angle potential parameter has been observed in the other linear CG models such as the Brannigan-Philips-Brown model^33^ and the Cooke-Kremer-Deserno model.^34^ The density profile (Fig. 3) also showed good agreement with the mapped atomistic results of a POPC bilayer.

**Table 2.** Physical properties of POPC bilayers. Standard errors are given in parentheses.

|  | CG |  | AA |
| --- | --- | --- | --- |
| Angle parameter | $0.5 k_B T$ | $2.0 k_B T$ | |
| APL [$\text{\AA}^2$] | 66.2 (0.0) | 59.7 (0.1) | 65.4 (0.2) |
| $K_A$ [$\text{mN/m}$] | 151.7 (4.4) | 224.3 (8.3) | 221.4 (16.3) |
| $K_C$ [$k_B T$] | 16.6 (0.3) | 35.7 (1.0) | 31.1 (2.3)<sup>a)</sup> |
| O.P.1 | 0.76 (0.02) | 0.80 (0.01) | 0.75 (0.01) |
| O.P.2 | 0.70 (0.02) | 0.75 (0.02) | 0.75 (0.02) |

a) Values from Ref. 12

**FIGURE 3.**
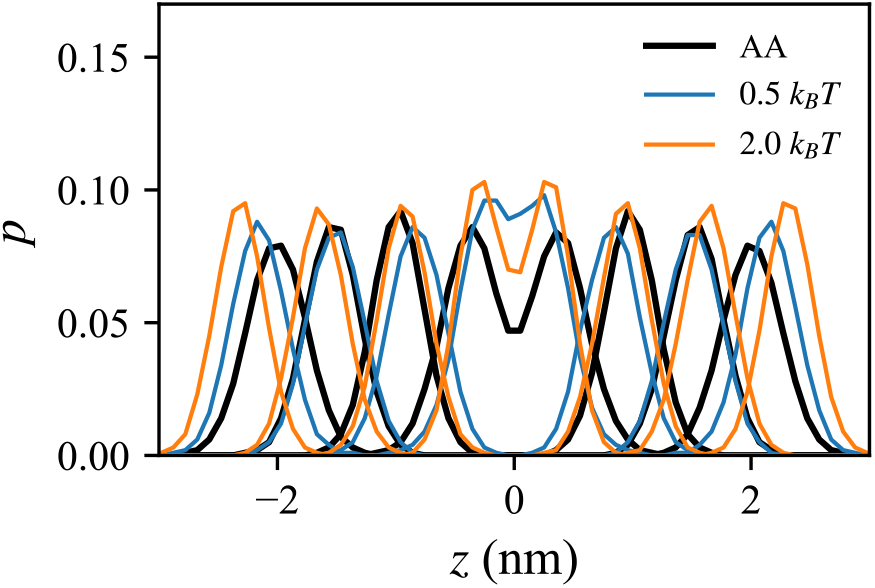
Comparison of the number distribution in the bilayer normal between the mapped AA trajectory (black) and the CG simulation with an angle parameter of 0.5 *k*_B_*T* (blue) or 2.0 *k*_B_*T* (orange).

A linear, 4-bead TG model was derived from the PL model (Figs. 1 and 4a). We characterized the properties of a bulk TG system containing 216 TG molecules. The volume of each TG molecule and the interfacial tension at the water interface were 1.30 ± 0.0 nm^3^ and 26.1 ± 0.3 mN/m, comparable to experimental data of those quantities,^35–36^ with 1.64 nm^3^ and 32 mN/m, respectively. Our model presented here is purely phenomenological and is not directly linked to any underlying AA systems. A discussion on CG representability and transferability issues can be found in Ref. 37.

**FIGURE 4.**
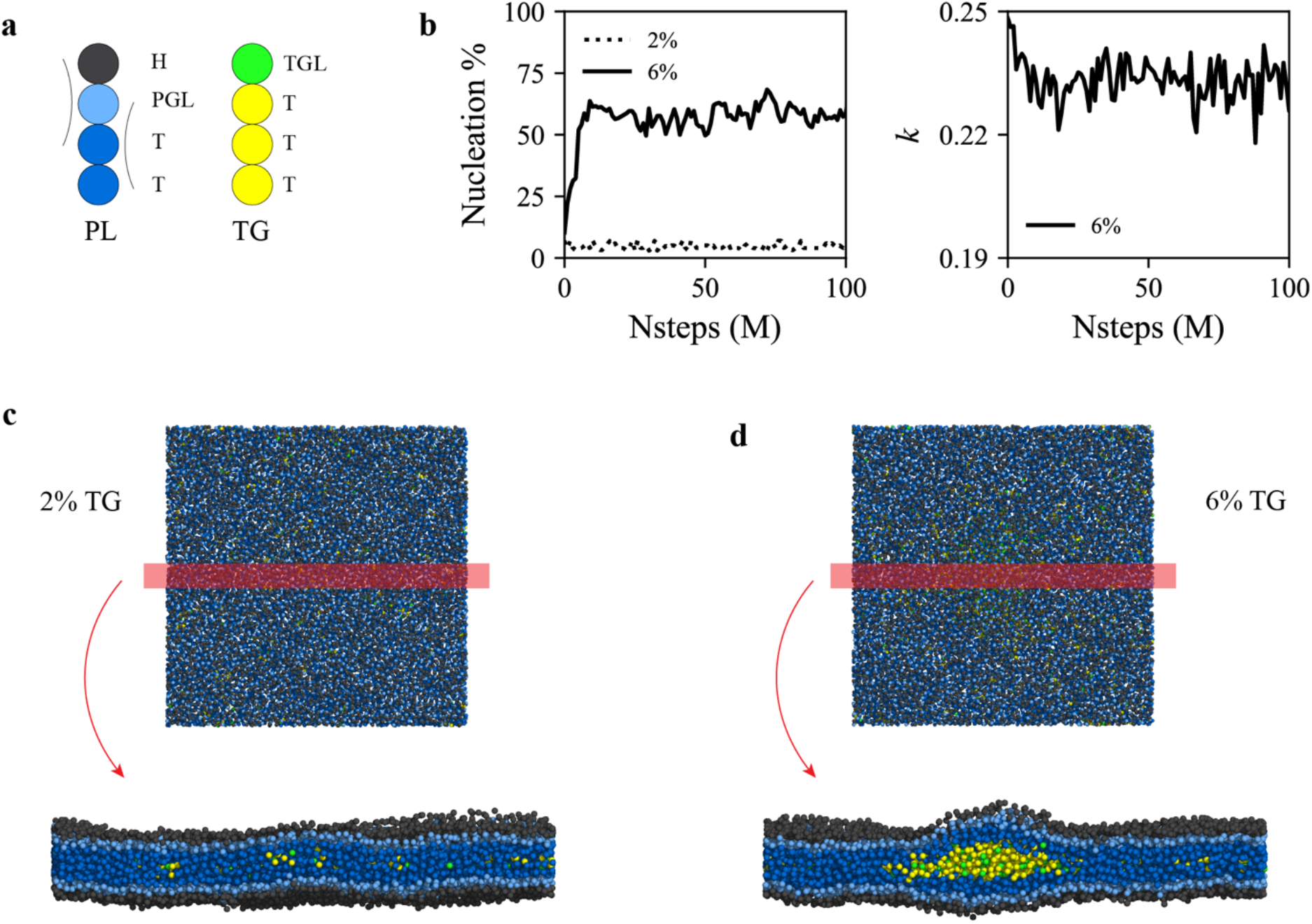
TG concentration-dependent nucleation. (a) Illustration of the PL and TG models used in this study. Each CG type is written next to each CG bead. Arcs represent harmonic angle potentials. The same color code is used in the rest of the study. (b) Nucleation % (left) and anisotropy (right) for bilayers containing 2% mol (dotted) or 6% mol (solid) TG. For visual clarity, anisotropy of the 2% mol TG bilayer is not shown. The last snapshots (100 M MD time steps) of the bilayers containing (c) 2% mol or (d) 6% mol TG are shown. Simulations discussed here were run with an angle potential parameter of 2.0 *k*_B_*T*.

### TG concentration-dependent nucleation

Experiments measuring the solubility of triolein in a POPC bilayer reported that ~ 2.4% mol TG can be dispersed in the membrane before phase nucleation occurs.^38^ If bilayers contain fewer TG molecules than this critical concentration, TG is dissolved in PL, whereas TG forms a distinct phase at the critical concentration or above. To model the TG concentration-dependent nucleation behavior, we simulated bilayers with two different TG concentrations, 2% mol and 6% mol TG. Consistent with the experimental data, TG underwent nucleation at 6% mol (Fig. 4b). In this case, the TG blister had anisotropy close to 0.25, representing a flat structure, to minimize membrane deformation (Figs. 4b and 4d). However, for TG concentration of 2% mol, TG did not nucleate but remained dissolved in the PL phase. (Figs. 4b and 4c). The simulations discussed here were run with an angle potential parameter of 2.0 *k*_B_*T*. In the bilayer simulations with an angle potential parameter of 0.5 *k*_B_*T*, we also observed TG dissolution at the 2% mol TG bilayer and TG nucleation at the 6% mol TG bilayer. By evaluating the nucleation PMF at those two different concentrations and with two different angle parameters, we will later discuss how TG concentrations and angle parameters change the free energy minimum and morphology of TG lenses.

### Ostwald ripening and PL rigidity-dependent lens shape

To study the mechanism of LD growth and PL rigidity-dependent lens shapes, a spherical bilayer membrane containing 6% mol TG with a diameter of 40 nm was simulated. We performed CG MD simulations with two different angle potential parameters, 0.5 *k*_B_*T* and 2.0 *k*_B_*T*. In both cases, we found that the final structure had one large TG lens between the PL leaflets. Examination of the simulations revealed two distinct mechanisms of LD growth, Ostwald ripening and coalescence of oil phases. In the simulation with stiff PLs, Ostwald ripening was observed (Figs. 5a and 5b). Two principal TG lenses were generated, and both lenses grew by attracting the neighboring TG molecules up to 30 M MD time steps. However, after that, the smaller cluster shrank and eventually dissolved, while the larger cluster grew in size. Since the two oil lenses were far apart, this process was not due to oil coalescence but due to Ostwald ripening. It should be noted that Ostwald ripening of LDs was recently seen experimentally.^39^ In contrast, simulations with soft PLs showed coalescing of clusters as indicated by a sharp increase of the TG number in the largest cluster and a sharp decrease in the second largest cluster. We also simulated a vesicle containing a 1:1 ratio mixture of stiff and soft PLs. In this case, the largest and the third largest oil lenses merged, while the second largest TG cluster gradually disappeared by Ostwald ripening (Fig. 6). However, we did not observe any sorting of PLs, and both stiff and soft PLs were equally distributed in the bilayer.

**FIGURE 5.**
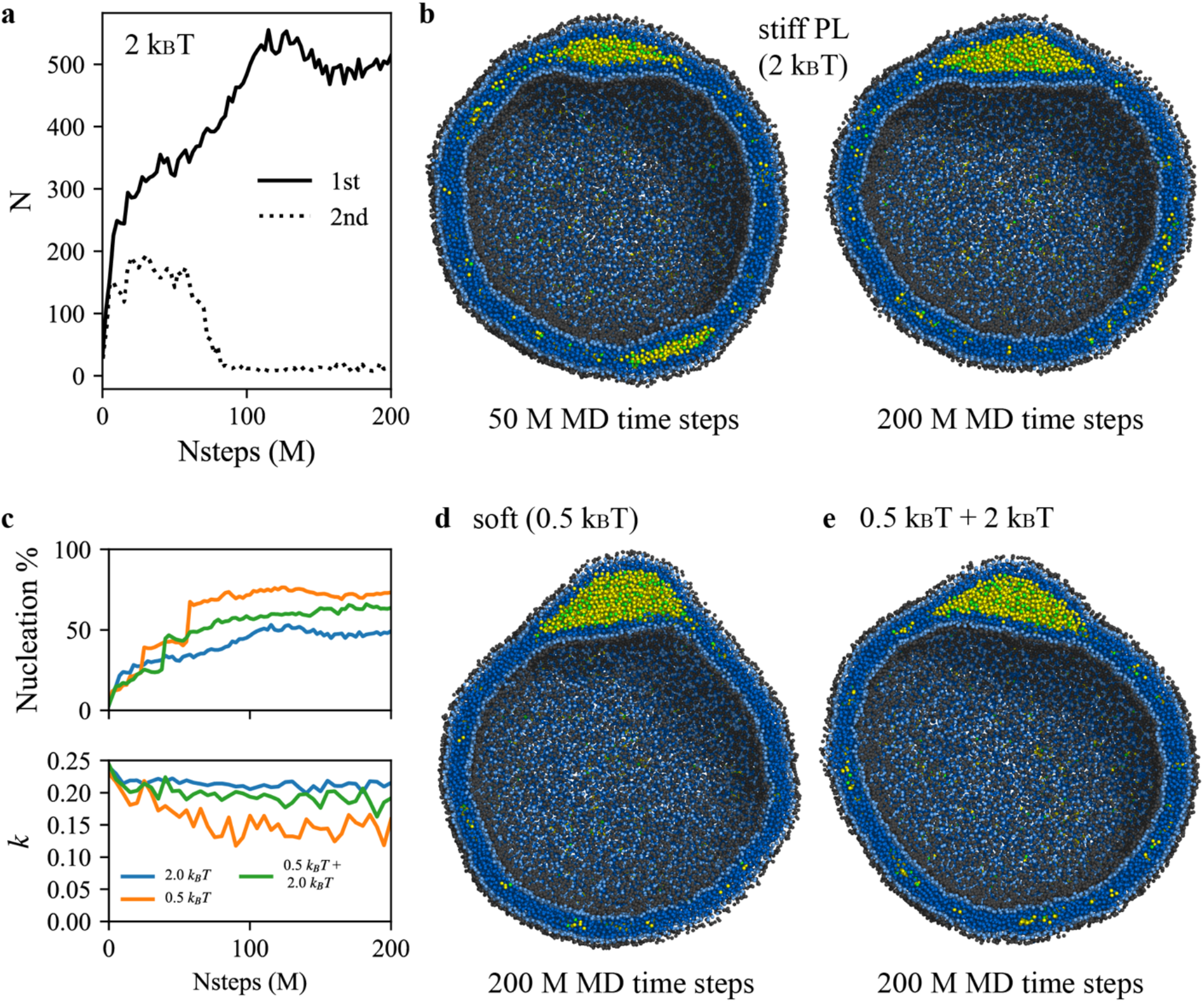
Ostwald ripening and PL rigidity-dependent LD shape. (a) The number of TG in the first (solid) and second (dotted) largest cluster. Simulations were run with an angle parameter of 2 *k*_B_*T*. (b) The interior view of the simulation at 50 M MD time steps (left) and 200 M MD time steps (right). (c) Nucleation % and anisotropy with simulation times. The first (blue) and second (orange) systems have angle parameters of 2 *k*_B_*T* and 0.5 *k*_B_*T*, respectively. The third system (green) consists of a 1:1 ratio mixture between stiff and soft PLs. The interior view at 200 M MD time steps of the (d) second and (e) third system.

**FIGURE 6.**
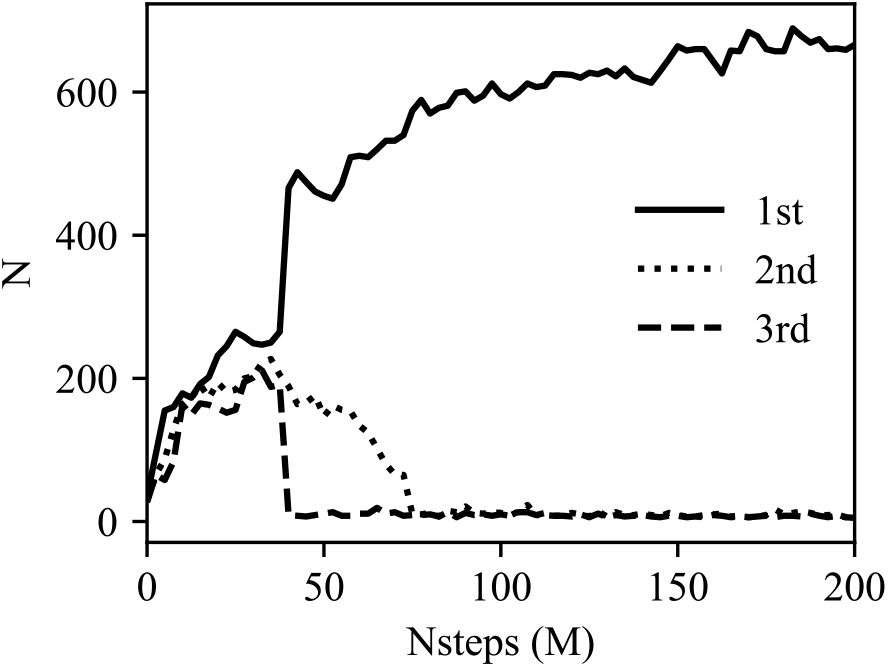
Oil coalescence and Ostwald ripening. Vesicular system consisting of a 1:1 ratio mixture between soft and stiff PLs. The number of TG molecules in the first (solid line), second (dotted line), and third largest cluster (dashed line). Merging between the first and third largest clusters happens at 45 M MD time steps, indicated by a sharp increase or decrease in the number of TG. The second cluster gradually disappears by Ostwald ripening.

How does membrane rigidity impact the shape of oil lens? We found significant differences in anisotropy and the shape of a TG cluster in the vesicular simulations. While the system with stiff PLs showed a flat oil blister, characterized by the high anisotropy, the system with soft PLs had a spherical oil blister with the low anisotropy (Figs. 5b, 5c, and 5d). Given the same bulk energy per volume, this can be understood as a TG’s response to a high membrane deformation penalty due to the PL’s high rigidity. We also observed that PL rigidity changed the nucleation percentage (Fig. 5c). With soft PLs, the equilibrated nucleation percentage increased. Interestingly, a vesicle that contained a 1:1 ratio of soft and stiff PLs presented the nucleation percentage and anisotropy values between those in single-component PL vesicles.

### Asymmetric tension and budding

Two spherical systems with a diameter of 40 nm and an angle parameter of 2.0 *k*_B_*T* were set up by varying the number of PLs in the inner leaflet while fixing the number of PLs in the outer leaflet. With this approach we imposed asymmetric tension between the monolayers. Consistent with the recent experimental studies,^40–41^ we confirmed asymmetric PL density controls the budding direction (Fig. 7). A TG blister budded toward the leaflet that better offers the PL coverage of TG. In the first simulation, where the ratio of the number of PLs in the inner leaflet to that in the outer leaflet is 0.69, a TG lens budded to the outer leaflet representing the cytosolic side of an ER membrane. In the other simulation, where the ratio is 0.89, the budding direction was reversed into the inside of the vesicle, representing the ER lumen. In contrast, a TG lens remained in a bilayer membrane in the simulation where the ratio is 0.79.

**FIGURE 7.**
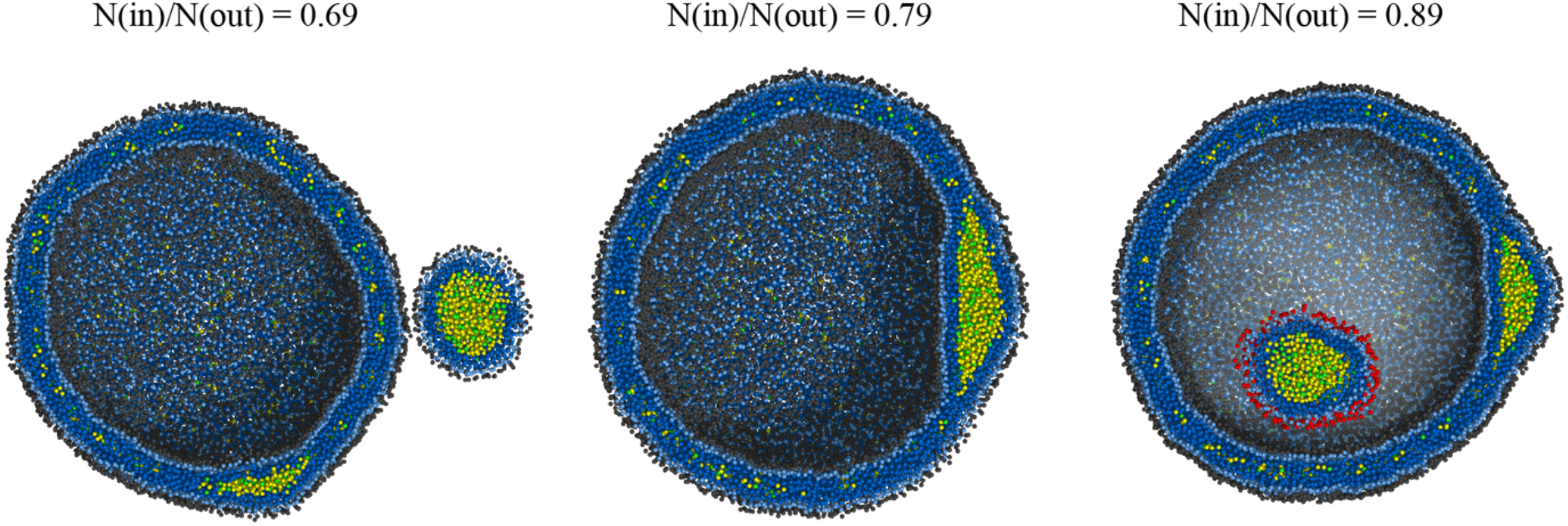
PL density-dependent budding. LDs bud toward the monolayer that exhibits better PL coverage of a TG cluster. The initial ratio of the PL number in the inner leaflet to that in the outer leaflet is shown. For visual clarity, the PL head group of the budded LD is shown in red in the right figure. Simulations were run with an angle potential parameter of 2.0 *k*_B_*T*.

### TG nucleation PMF

To estimate the free energy of TG nucleation as a function of the degree of nucleation and anisotropy, we carried out well-tempered metadynamics simulations as described earlier in Methods. Because the degree of nucleation (or equivalently the nucleation percentage) is not a continuous function and therefore cannot be biased, we instead biased the sum of the coordination number of the TGL atoms in the largest cluster. A high correlation between the degree of nucleation and the sum of the coordination number in the largest cluster was achieved (Fig. 8 top and second panels from top) by using a sharp switching function (Fig. 2). However, if one uses a broad switching function, a high correlation between the nucleation percentage and the coordination number is not guaranteed. A broad switching function can result in formation of several clusters, each having a dense aggregation of TG, which is not preferred.

**FIGURE 8.**
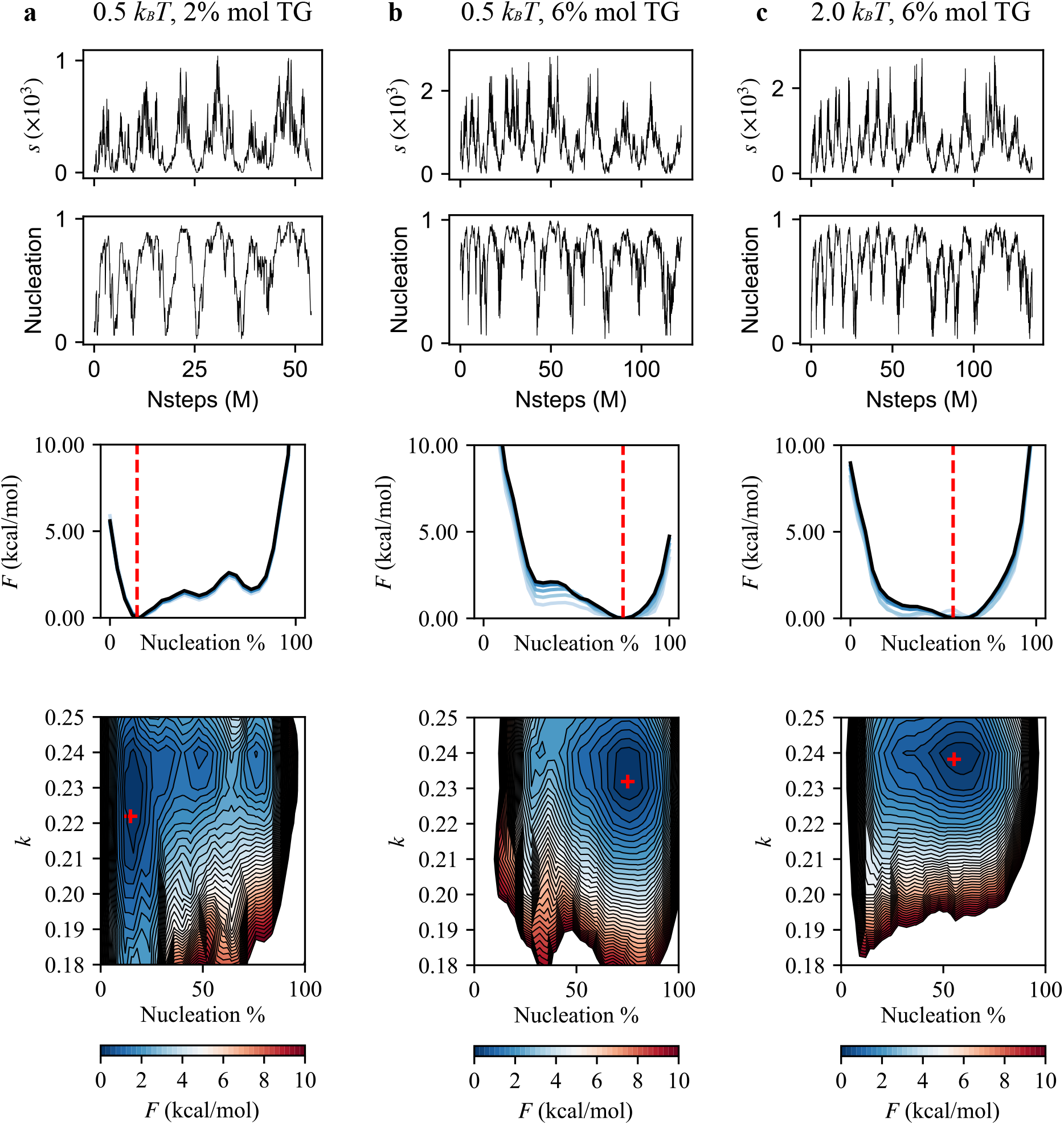
TG nucleation free energy. From top to bottom: (first row) the biased collective variable, the sum of the coordination number of TGL atoms in the largest cluster, in the biased trajectory. (second row) The degree of nucleation calculated in the biased trajectory. (third row) The calculated PMF as a function of nucleation % by reweighting the biased simulation. The red dashed line indicates the equilibrated nucleation % value that was obtained from the unbiased simulation of the same bilayer. The blue lines indicate how PMFs evolve with simulation times to show convergence. The lighter the color, the less simulation frames were used for calculating the PMF. (fourth row) The calculated PMF as a function of nucleation % and anisotropy. The red marker indicates the equilibrated values of nucleation % and anisotropy from the unbiased simulation of the same bilayer. (a) The simulation was run with an angle parameter of 0.5 *k*_B_*T* and contains 2% mol TG. (b) Angle parameter of 0.5 *k*_B_*T* in the 6% mol TG bilayer. (c) Angle parameter of 2.0 *k*_B_*T* in the 6% mol TG bilayer.

Small bilayers containing 2% mol and 6% mol TG were simulated for these calculations. In the case of 6% mol TG, two different angle potential parameters, 0.5 *k*_B_*T* and 2.0 *k*_B_*T*, were used. The coordination number (Fig. 8 top panels) sampled most of the region several times, except the very low and high values. Accordingly, the degree of TG nucleation was widely sampled in the biased simulations (Fig. 8 second panels from top). We reweighted the trajectory to calculate the TG nucleation PMF as a function of TG nucleation percentage and anisotropy (see Methods).

We first compared the results of the biased trajectories with those of the unbiased trajectories. Consistent with the unbiased trajectory that did not have TG nucleation in a bilayer containing 2% mol TG, the PMF of the same bilayer indicated a free energy minimum at the low nucleation percentage. The PMF also increased with the nucleation percentage (Fig. 8a). In contrast, the bilayer containing 6% mol TG had a free energy minimum at the high nucleation %. Furthermore, denoted by red lines (Fig. 8 third panels from top) and red markers (Fig. 8 bottom panels), the equilibrated values of the nucleation percentage and anisotropy from the unbiased simulations agreed well with the free energy minima in the calculated PMFs. Overall, our PMFs showed good agreement with the unbiased trajectories, therefore confirming the robustness and convergence of our biased simulations.

We next investigated the influence of the angle potential parameter. Soft PLs enhanced the nucleation percentage and decreased anisotropy, which is confirmed in our PMFs by comparing the free energy minima (Fig. 8b and 8c). The bilayer with an angle parameter of 0.5 *k*_B_*T* had an increased nucleation percentage and decreased anisotropy compared to the bilayer with an angle parameter of 2.0 *k*_B_*T*. Also, we observed that the contour of 10 kcal/mol became more extended toward the low anisotropy region in the simulation with soft PLs than the simulation with stiff PLs.

Finally, we compared TG nucleation with argon gas nucleation. In argon gas nucleation, the whole range of anisotropy from 0 to 0.25 is sampled,^30^ while in TG nucleation, anisotropy is only limited to the high values. This suggests different thermodynamic features rule nucleation. Argon gas nucleation has the surface tension energy penalty, which makes a cluster tend to become spherical. In TG nucleation, membrane deformation rather than surface tension works against TG nucleation, which makes a TG blister sample only the planar region (high anisotropy region). However, one might expect that the lower anisotropy region will be more sampled when a system becomes bigger.

## DISCUSSION

In this study, the biophysics of LD emergence was investigated with a highly tunable, phenomenological CG model.^13^ We characterized the physical properties of a 4-bead PL model (head - glycerol - tail - tail), including the APL, bilayer area compressibility, bending modulus, order parameters, and density profile. Although we used a generic model, the CG results compared well with the AA results of a POPC bilayer membrane, which is the primary component of PLs in a mammalian cell.^42^ Two spring constants of harmonic angles, 0.5 *k*_B_*T* and 2.0 *k*_B_*T*, were used to control PL rigidity. Based on the pair potentials between PL atoms, we made a 4-bead TG model (glycerol - tail - tail - tail) that showed the concentration-dependent behavior. If the TG concentration is above the critical, TG nucleates a lens.

LD biogenesis is driven by neutral lipids’ bulk energy and is opposed by surface tension and membrane deformation energy. When LDs are small, the membrane deformation energy is dominant. However, as the LDs grow, the surface tension energy takes over as it is proportional to the surface area. A characteristic length of this transition is predicted to be 10 nm – 20 nm.^8–9^ In our simulations, the LDs are smaller than this characteristic length, therefore we mostly demonstrate the interplay between TG lensing and membrane deformation energy, in which the latter was controlled via PL rigidity.

In the simulations of a vesicle with a diameter of 40 nm, we showed PL density-dependent budding phenomena, which is consistent with the recent experimental papers.^40–41^ In the simulation where there was an excess PL in the inner leaflet, LDs budded to the center of the sphere, whereas LDs budded to the outside of the sphere when there was an excess PL in the outer leaflet. Therefore, in a closed, tunable-sized bilayer system, the balance of PLs between the inner and outer leaflets determines the budding directionality. In cells, the ER bilayer is an open system due to vast amounts of ER, scramblases or flippases,^43^ and de novo PL synthesis; Yet, there are still apolipoprotein B-free lumenal LDs.^44–45^ Using our model, a possible explanation could be the local accumulation of excess PLs in the lumenal leaflet or stiff PLs in the cytosolic leaflet.

In the vesicular simulations that do not show budding, two mechanisms contribute to the formation of one large TG lens between the PL leaflets: Lens coalescence and Ostwald ripening. Ostwald ripening was shown in our trajectories where only one of two distanced TG lenses grew and the other became gradually dissolved. The experimental evidence of Ostwald ripening of LDs was reported in Ref. 39. Although not directly related to lens coalescence in a bilayer membrane, Fsp27-mediated LD coalescence at the LD contact site was shown in Ref. 46.

We also studied the correlation between PL rigidity and the shape of a TG lenses. With reduced PL rigidity, LDs became more spherical, and the nucleation percentage increased. To support our conclusions, we calculated the TG nucleation PMF in a flat bilayer. By biasing the sum of the coordination number of TG glycerol (TGL) atoms in the largest cluster and reweighting the biased trajectory, we computed the PMF as a function of the nucleation percentage and anisotropy. Consistent with the vesicular simulations, the free energy minimum is located at the lower anisotropy and higher nucleation percentage with reduced PL rigidity.

Finally, we note that TG itself can significantly alter the membrane properties. It was recently shown experimentally^47^ and computationally^12^ that TG reduces bending modulus of a bilayer membrane. The other neutral lipid, diacylglycerol, can also reduce membrane rigidity.^12, 48^

## CONCLUSIONS

In a bilayer membrane, TG nucleation is driven by its bulk energy. However, membrane deformation incurs an energy penalty on nucleation at the initial phases of LD formation. Our CG simulations demonstrated the competing effects of TG lensing and membrane deformation. We showed high membrane rigidity reduces the nucleation percentage and increases anisotropy of a TG lens, confirmed in large-scale vesicular simulations and the calculations of the TG nucleation free energy. In addition, two distinct mechanisms that govern LD growth, Ostwald ripening and coalescence of oil lenses, were shown in vesicular simulations. Finally, the LD budding direction was controlled by the number of PLs in the inner and outer leaflets. Taken together, we provide a better understanding of LD formation at the initial steps, validate the reported experiments, and conclude that membrane rigidity serves as a key factor in the formation and shape of a TG lens.

## SUPPORTING INFORMATION

The PLUMED script for biasing simulations is submitted along with the manuscript.

## AUTHOR CONTRIBUTIONS

S.K. and G.A.V. designed the research. S.K. performed the simulations. All authors analyzed the results and wrote the study.

## ACKNOWLEDGEMENTS

This research was supported by a grant from the National Institute of General Medical Science (NIGMS) of the National Institutes of Health (NIH), grant R01-GM063796. The computer simulations were performed on the Midway2 and Midway3 clusters at the University of Chicago and the Bridges2 computer at the Pittsburgh Supercomputing Center (PSC) through allocation MCA94P017, with resources provided by the Extreme Science and Engineering Discovery Environment (XSEDE) supported by NSF grant ACI-1548562. We thank Federico Giberti, Jeeyun Chung, and Alex Pak for their valuable perspectives and discussion.

---

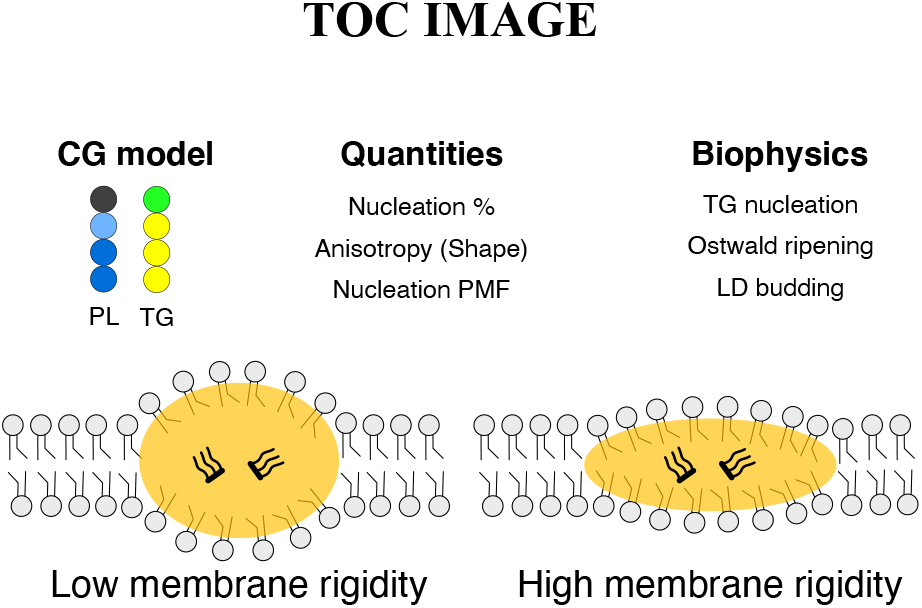

