## Supplemental File for "Key Factors Governing Initial Stages of Lipid Droplet Formation"

### PLUMED script for biasing simulations

```
MOLINFO STRUCTURE=molinfo.pdb

lq:     COORDINATIONNUMBER  SPECIES={@mda:{name TGI}}  SWITCH={CUBIC D_0=1.95 D_MAX=2.0}  LOWMEM
cm:     CONTACT_MATRIX      ATOMS=lq                   SWITCH={CUBIC D_0=1.95 D_MAX=2.0}
dfs:    DFSCLUSTERING       MATRIX=cm                  LOWMEM
clust1: CLUSTER_PROPERTIES  CLUSTERS=dfs   CLUSTER=1   SUM

METAD ...
label=m
ARG=clust1.sum
HEIGHT=2.0
SIGMA=10
PACE=500
GRID_MIN=0
GRID_MAX=3000
GRID_WSTRIDE=500000
GRID_WFILE=grid.dat
TEMP=310
BIASFACTOR=50
CALC_RCT
... METAD

ss: CLUSTER_NATOMS CLUSTERS=dfs CLUSTER=1
PRINT ARG=clust1.sum,ss,m.bias,m.rbias STRIDE=500 FILE=colvar
```
